# Gray Matter Differentiation Between Two Subordinate Personality Traits of Extraversion: Enthusiasm and Assertiveness

**DOI:** 10.1101/2021.11.08.467744

**Authors:** Magnus Frisk, Gránit Kastrati, Jörgen Rosén, Kristoffer N. T. Månsson, Fredrik Åhs

## Abstract

Enthusiasm and assertiveness are two subordinate personality traits of extraversion. These traits reflect different aspects of extroversion and have distinct implications on mental health. Whereas enthusiasm predicts satisfaction in life and positive relationships, assertiveness predicts psychological distress and reduced social support. The neural basis of these subordinate traits is not well understood. To investigate brain regions where enthusiasm and assertiveness have diverging relationship with morphology, enthusiasm and assertiveness were regressed to gray matter volume (GMV) across the whole brain in a sample of 301 healthy individuals. A significant interaction was found between enthusiasm and assertiveness in the left angular gyrus (*t*(296) = 4.18, family wise error corrected, FWE *p* = .001 (cluster-level); Cluster size *=* 880 voxels). Larger GMV in this area was associated with more enthusiasm and less assertiveness. Our study emphasizes the value of separating extraversion into its subordinate traits when investigating associations to neuroanatomy.

## 1. Introduction

Extraversion has previously been split into a dyad (Deyoung, Quilty, & Peterson, 2007; Watson & Clark, 1997). This dyad has been labelled *Enthusiasm* and *assertiveness* (Deyoung et al., 2007). Enthusiasm includes traits such as gregariousness, friendliness, and enjoyment of rewards, while assertiveness includes agency, dominance, and the desire to seek out rewards (Deyoung et al., 2007; Deyoung, 2010). Both traits encompass behavioral tendencies in social interactions. However, each trait may likely influence social relationships distinctively. For instance, assertiveness has been associated with reduced social support while enthusiasm is associated with more positive relationships (Newby-Fraser & Schlebusch, 1997; Sun, Kaufman, & Smillie, 2018). Furthermore, enthusiasm has been reported to mitigate psychopathology while assertiveness sometimes shows an increased risk for psychiatric disorders. Enthusiasm is associated with higher life satisfaction and fewer negative emotions whereas assertiveness is associated with more negative emotions, externalizing psychopathology, and substance use disorder (Sun, Kaufman, & Smillie, 2018; Walton, Pantoja, & McDermut, 2018; Watson, Stasik, Ellickson-larew, & Stanton, 2015; Watson et al., 2019). Considering that implications associated with mental health differ substantially between enthusiasm and assertiveness, it would be elucidating to determine how these traits are associated with brain morphology.

Past research has rigorously studied the association between extraversion and gray matter volume (GMV) in the human brain (Lai et al., 2019; Wacker & Smillie, 2015). However, heterogeneous results between studies have made it difficult to ascertain any reliable associations between extraversion and brain morphology. One limitation in past research may be attributed to the use of small samples (Button et al., 2013; Kharabian Masouleh, Eickhoff, Hoffstaedter, Genon, 2019). Another issue could be the complexity of extraversion. Possibly, correlations between extraversion and GMV could be obscured if one subordinate trait of extraversion is positively correlated, while another trait is negatively correlated. Deyoung (2010) suggested that by decomposing extraversion into subordinate traits, associations to more specific regions of the brain may be revealed. The idea of examining subordinate traits of extraversion has shown some promise with at least one smaller study (83 participants) that has identified a few brain regions associated with subordinate traits of extraversion (Grodin & White, 2015). These regions consisted of the medial orbitofrontal cortex, parahippocampal gyrus, left cingulate gyrus, and left precentral gyrus. However, a more recent study with a much larger sample of 1107 participants, did not find any significant associations between subordinate traits of extraversion (operationalized as warmth, gregariousness, assertiveness, activity, excitement-seeking, and positive emotions) and brain morphology (Avinun, Israel, Knodt, & Hariri, 2020). Avinun et al. (2020) used non-target personality traits and sex as covariates and utilized multiple parcellation schemes (cortical thickness and surface area, subcortical volume, and white matter microstructural integrity). However, their study did not investigate interactions between subordinate traits of personality and brain morphology. Considering that prior research has shown that assertiveness and enthusiasm have distinct associations with mental health, such interactions may be of interest to better understand psychopathology.

While previous studies have investigated subordinate traits of extraversion independently from each other, this study primarily focused on brain regions where the relationship between GMV and enthusiasm where different from the relationship between GMV and assertiveness. The purpose of this approach was to locate brain areas where morphological development and associated cognitive functionalities are differentiated between enthusiasm and assertiveness. Importantly, regions where enthusiasm and assertiveness differ may further explain how come these traits are idiosyncratically associated with mental health. To this end, the present study investigated interactions between subordinate traits of extraversion and GMV across the whole brain in a sample of 301 individuals.

## 2. Experimental Procedures

### 2.1 Participants and Procedure

Anatomical brain scans were obtained from a research project studying genetic influences on brain function in twins. Here, correlations between brain morphology and subordinate traits of extraversion were evaluated without considering genetic influences as these results will be reported elsewhere. 301 Adults (179 female/122 male) were recruited through the Swedish Twin Registry (Zagai, Lichtenstein, Pedersen & Magnusson, 2019). Mean age of participants was 33.84 ± 0.49 and ages ranged from 20 to 58. Two-hundred-fifty-eight of the participants reported being right-handed. Exclusion criteria were having any implants preventing MRI scanning, ongoing pregnancy or treatment with psychotropic medication, claustrophobia, drug or alcohol related problems or past surgical procedures of either heart or brain. Participants were interviewed over the phone and then invited to the MR Research Center in Solna, Sweden, for MR-scanning. The study was approved by the Uppsala Ethical Review Board and all participants signed informed consent. Personality data was collected with an online questionnaire using LimeSurvey (www.limesurvey.org), which participants completed either before or after MR-scanning. Each participant received 1000 SEK (∼$100) for participating in the study.

### 2.3 Extraversion

The Swedish version of Big Five Inventory (BFI) was used to assess different traits of extraversion. Measurement properties and internal consistency of the Swedish BFI have been validated, with the extraversion related items presenting a Cronbach’s alpha of, α = .84 (Zakrisson, 2010). Each item in BFI has five response options: disagree strongly = 1 point, disagree a little = 2 points, neither disagree nor agree = 3 points, agree a little = 4 points and agree strongly = 5 points. Participants responded to the following items measuring extraversion: (1) I see myself as someone who is talkative, (2) I see myself as someone who is reserved (reversed item), (3) I see myself as someone who is full of energy, (4) I see myself as someone who generates a lot of enthusiasm, (5) I see myself as someone who tends to be quiet (reversed item), (6) I see myself as someone who has an assertive personality, (7) I see myself as someone who is sometimes shy, inhibited (reversed item), and (8) I see myself as someone who is outgoing, sociable. Responses to reversed item were changed to make all items positively loaded to extraversion.

### 2.4 Magnetic Resonance Imaging

Magnetic Resonance Imaging was conducted using a General Electric 750 (General Electrics, Milwaukee, WI, USA; Karolinska Institutet, 2015) 3T MRI scanner. Anatomical whole brain T1-weighted images were obtained from all participants while they were resting with eyes closed. During scanning, a radio-frequency pulse was initialized every 7.9 s (TR). After each pulse, the highest feedback signal was picked up after 2.8 ms (TE). Each pulse sent from the scanner had an angle of 11 degrees to the surrounding magnetic field (flip angle). Voxel size was 0.9735 mm × 0.9735 mm × 1.000 mm.

### 2.5 Gray Matter Volume

Preprocessing and analysis of images were performed using SPM12 (Version 7771, https://www.fil.ion.ucl.ac.uk/spm/) and CAT12 (Version 1318, https://www.neuro.uni-jena.de/cat/). Images were segmented and then spatially registered using Geodesic Shooting in CAT12. Spatial registration aligns images to minimize global discrepancies of displacement. Normalization to a common voxel-space is used to facilitate interpretation of results and enable inter-study comparisons. In this study, affine regularization used the ICBM-space template for European brains (Template_0_IXI555_MNI152) when normalizing to MNI space. Following the completion of image pre-processing, the voxel size was resampled to 1.5 mm × 1.5 mm × 1.5 mm. An Isotropic Gaussian Kernel of 8 mm full at half maximum was used to smooth images. Following statistical analysis, MNI coordinates were anatomically labeled using Automated Anatomical Labelling Atlas 3 (Rolls, Huang, Lin, Feng, & Joliot, 2020).

### 2.6 Statistical Analyses

The correlational structure of extraversion was assessed using a principal component analysis in SPSS statistics (Version 25.0.0.2). Two components emerged with eigenvalues above 1 (Table 1) and were labelled enthusiasm (*eigenvalue = 4*.*56*) and assertiveness (*eigenvalue = 1*.*02*). All items excluding (6), I see myself as someone who has an assertive personality, loaded onto enthusiasm, whereas assertiveness was predominantly constituted by item (6). For each component and participant, a factor score was calculated from the factor loadings. These two scores were then used as predictor variables in regression analyses with GMV as dependent variable in SPM12. An explicit brain mask was used to restrict the analysis to voxels within the brain. Age and sex were included as covariates of no interest. Global nuisance effects were adjusted for by dividing each image with its global mean voxel value using the proportional scaling function in SPM12. A threshold of *p* =.001 (uncorrected) was applied and voxels surviving a family wise error corrected *p*-value (*pFWE*) of *p* < .05 at the cluster level are reported. To evaluate the magnitude and the direction of the association between GMV and 1) enthusiasm, and 2) assertiveness, another regression analysis was conducted in SPSS statistics using enthusiasm and assertiveness as predictor variables, age and sex as control variables, and eigenvalues of identified cluster as dependent variable.

**Table 1.**
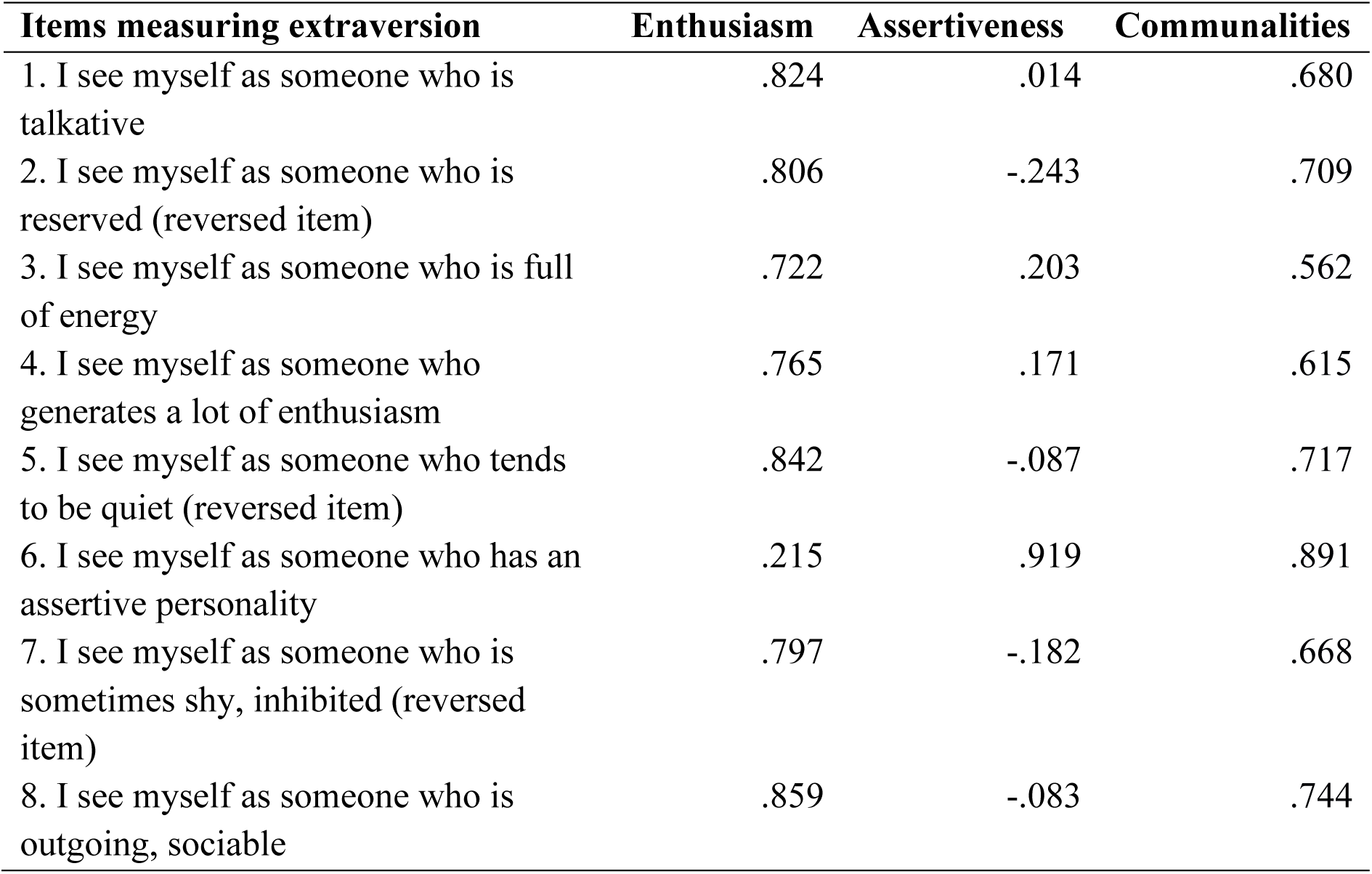
Factor loadings from Principal Component Analysis of extroversion items from the Swedish version of Big Five Inventory

## 3. Results

We found an interaction between the two subordinate traits of extroversion, enthusiasm and assertiveness, and GMV in a cluster of 880 voxels (1320 mm^3^) in the region of left angular gyrus (*t*(296) *=* 4.18; cluster-level *pFWE =* .001; MNI coordinates: x = -33, y = -72, z = 40). The interaction was explained by a positive correlation between enthusiasm and GMV, contrasted to a negative correlation between assertiveness and GMV. This result indicates that larger GMV is associated with more enthusiasm and less assertiveness. A scatter plot depicting enthusiasm and assertiveness plotted against GMV eigenvalues is shown in Figure 1. No significant associations were found between enthusiasm or assertiveness and GMV when analyzed separately at the level of the whole brain.

**Fig. 1.**
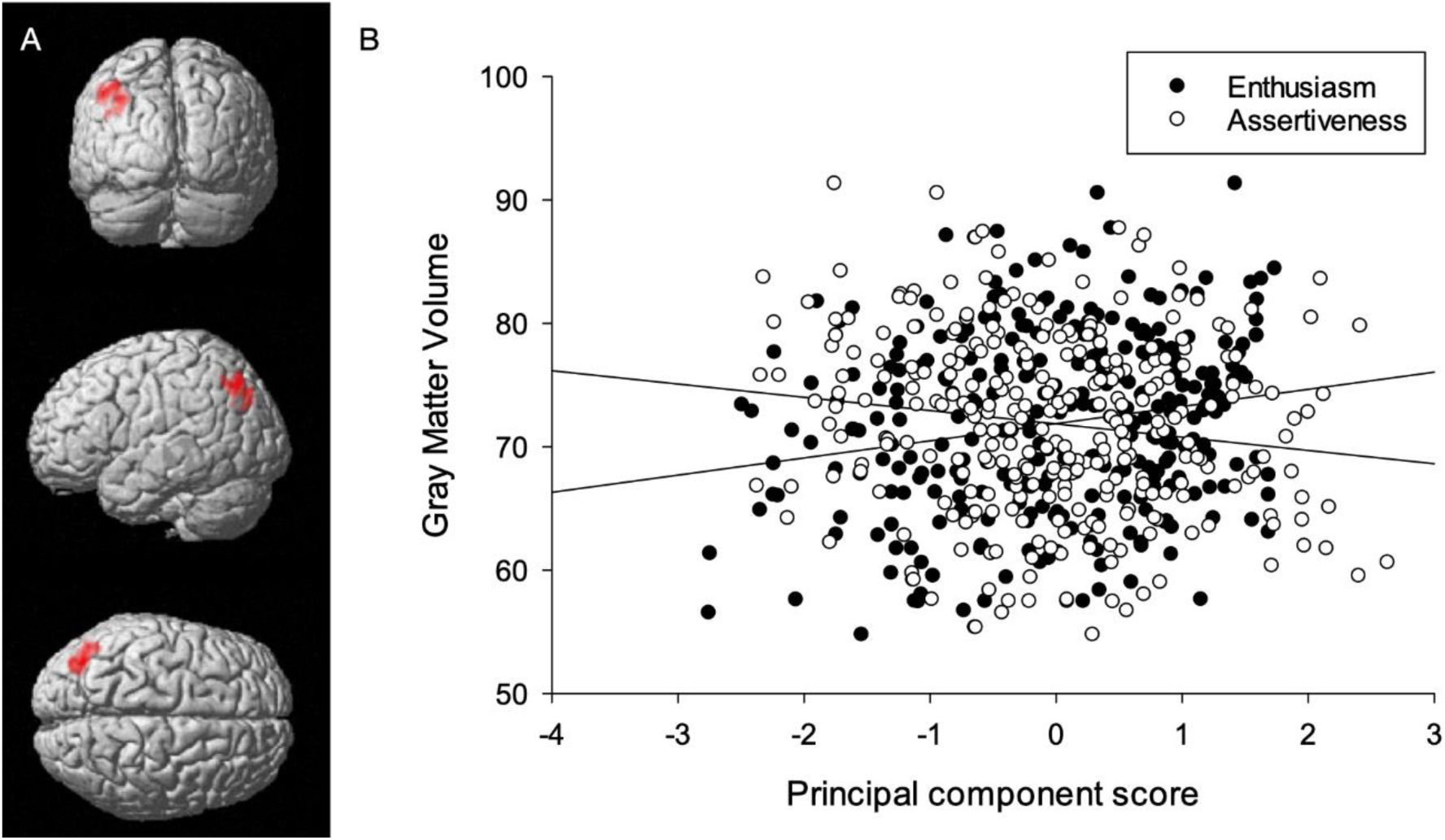
Interaction of enthusiasm and assertiveness with gray matter volume. A) Anatomical rendering of the brain showing a cluster of voxels in left angular gyrus where enthusiasm and assertiveness present opposing associations with GMV. B) Scatter plot showing a positive association between enthusiasm and GMV and a negative association between assertiveness and GMV.

A regression analysis (*r*^*2*^ = .088) with the extracted cluster eigenvalues as dependent variable yielded standardized beta coefficients of *β =* .22 *(p <* .001) for enthusiasm and *β* = -.18 (*p =* .002) for assertiveness. Age had a standardized beta coefficient of *β* = -.17 (*p* = .005) while sex had a non-significant contribution to the model.

## 4. Discussion

The purpose of the study was to evaluate if there are regions where two subordinate traits of extraversion, enthusiasm and assertiveness, differ from each other in their associations with GMV in the brain. Associations were tested through a whole brain voxel-wise regression analysis. For the left angular gyrus, we report a significant interaction between the subordinate traits of extraversion, enthusiasm and assertiveness, and GMV. This interaction was defined by a positive association between GMV and enthusiasm, and a negative association between GMV and assertiveness. Thus, individuals with greater GMV in the left angular gyrus, on average, rate themselves as more enthusiastic and less assertive.

### 4.1 Comparison to previous research

The main finding in our study may be explained by investigating the interaction of subordinate traits of extraversion and GMV. To our knowledge, no previous research has sought to identify brain regions where subordinate traits of extraversion are differentiated in their associations to GMV. When considering enthusiasm and assertiveness independently from each other, no significant associations to GMV were identified surviving correction for multiple comparisons at the level of the whole brain. This null result from our study is consistent with the most extensive study to date which investigated associations between subordinate traits of extraversion and brain morphology (Avinum et al., 2020). Our finding of a small negative association between age and GMV in left angular gyrus is likely best explained by the established consensus of a general reduction of gray matter in the process of aging.

### 4.2 Theory of mind as a possible common pathway for enthusiasm and assertiveness

Integrating information from large numbers of functional neuroimaging experiments, the angular gyrus emerges as a cross-modal hub important for the comprehension of events and complex social interaction (Seghier, 2013). Higher GMV in left angular gyrus has previously been associated with more accurate inferences of other people’s intentions and emotions (Laillier et al., 2019; Lewis et al., 2011). The positive association which we found between enthusiasm and GMV in left angular gyrus may thus be indicative of improved efficiency in theory of mind related abilities. Social interactions have shown to be one key factor in fostering the development of theory of mind during childhood (Derksen, Hunsche, Giroux, Connolly, & Bernstein, 2018). Possibly, enthusiasm may facilitate maintenance and continued development of theory of mind throughout adult life via endorsement of reciprocal social interactions. With assertiveness encompassing dominance in social interactions, this could have a different impact on angular gyrus related functionalities in comparison to enthusiasm. Input from others regarding emotions, thoughts, and mental states are all important in development of theory of mind (Ruffman, Slade, & Crowe, 2003; Slaughter, Peterson, & Mackintosh, 2007). Assertiveness may inhibit perception of these cues, as observational behavior is unlikely to occur simultaneous to assertion. With a reduced attention to surrounding social stimuli, top-down processes of angular gyrus are likely less stimulated. This contingency could be represented by the negative association which we found between assertiveness and GMV in left angular gyrus. It is worth mentioning that enhanced performance is sometimes associated with decreases in GMV, as part of neural maturity (Modroño et al., 2019). However, our analyses used age as a nuisance variable to control for maturity effects.

### 4.3 Implications for psychopathology

As enthusiasm and assertiveness are personality traits which both can be attributes within a single individual, interactive effects of these traits are important to consider. For instance, in addition to theory mind, assertiveness has shown to be beneficial in improving the quality of moral judgement (Hao & Liu, 2016; Kuramori, 1999). Possibly, assertiveness may enable reevaluation of beliefs when interacting with others. When misinterpreting feelings or intentions of others, asking questions may yield the opportunity to learn, improving the accuracy of mentalizing abilities over time. Thus, a moderate amount of assertiveness in combination with enthusiasm may be a balanced combination for social reciprocity, facilitating maturation of theory of mind. However, in cases where a high degree of assertiveness prevails in the absence of enthusiasm, maladaptive social strategies may instead influence the risk for psychopathology. For example, the mitigating effect enthusiasm has on psychopathology may at least partly be contingent with access to social support (Siedlecki, Salthouse, Oishi, & Jeswani, 2014). Conversely, with assertiveness having a depreciative effect on this contingency (Newby-Fraser & Schlebusch, 1997), the interaction of enthusiasm and assertiveness may either attenuate psychopathological outcomes, or drastically increase the risk. Lower enthusiasm and higher assertiveness may be a combination represented by low GMV in left angular gyrus, as suggested by our results. Indeed, GMV in the angular gyrus has been correlated with psychopathology across major depressive disorder, bipolar disorder and schizophrenia (Stein et al., 2021).

### 4.4 Recommendations for future research

Limited research has been conducted on the relationship between extraversion and theory of mind. From the few studies which we were able to find, some interesting results indicate that the lowest and highest percentiles of extraversion are associated with increased ability to determine emotional states in eye-gazes (King, 2015 Svenson & Guillen, 2020). What proportion of these finding may be attributed to enthusiasm or assertiveness remains unknown but marks an interesting topic for future research. We encourage continued research to investigate the association between subordinate traits of extraversion and theory of mind, as this may bring further clarity to the associations which we found in the region of left angular gyrus.

While discussing the potential relationships between the brain’s GMV and personality traits it is important to consider what the actual measure of GMV represent. Although it is commonly assumed that the brain’s GMV is relatively stable, the measure of GMV using T1-weighted images remains to be fully elucidated (e.g., Tardif et al. 2016). Some evidence shows that T1-weighted images are more sensitive than previously believed. For instance, Månsson and colleagues recently demonstrated activity-dependent changes in the brain’s GMV while individuals were engaged in a simple and passive visual task (Månsson et al., 2020). In our study, participants where at rest while undergoing T1-weighted imaging (i.e., no stimuli or task was present, thus reducing the risk for such confounds). Relative to the present results in this study, it could be that resting-state neural response in the left angular gyrus is associated with subordinate traits of extraversion. Nevertheless, it would be informative to also extend this research to measures of GMV during participants’ exposure to social stimuli.

## 5. Limitations

Our approach to obtain enthusiasm and assertiveness may entail some limitations when making comparisons to previous work. Rather than having utilized a personality assessment tool with a large subset of items, the Swedish Big Five Inventory only contained eight items. As a result, only one item had a clear loading on the factor which were labelled assertiveness (“I see myself as someone who has an assertive personality”). Since individual participants may have different views on what is an assertive personality, additional items describing assertive traits would have been beneficial to assure more reliable self-ratings. Furthermore, one critique of orthogonalizing enthusiasm and assertiveness through factor analysis may be that these traits have previously shown to have at least some correlation with each other (Deyoung, Weisberg, Quilty, & Peterson, 2013). This may be due to the concept of positive emotionality, which is sometimes shared between enthusiasm and assertiveness (Morrone-Strupinsky & Lane, 2007; Smillie, et al., 2015). Since our study used orthogonal representations of enthusiasm and assertiveness, the content of enthusiasm and assertiveness may not completely overlap with studies that have used the same labels for correlated representations.

Further, considering relatively recent critique towards cluster-extent thresholds (Eklund, Nichols, & Knutsson 2016), it is worth mentioning that the p-value of the identified cluster was pFWE = 0.001. It has been demonstrated that with a p-value of pFWE = 0.001 at the cluster level, results are reasonably robust (Kessler, Angstadt, & Sripada 2017). Hence, protection against false positives can be considered adequate in our study.

## 6. Conclusion

We found that two subordinate traits of extroversion, enthusiasm and assertiveness, differed in their associations with GMV in left angular gyrus. This interaction was represented by enthusiasm having a positive association with GMV whereas assertiveness displayed a negative relationship. Angular gyrus is a region which has consistently been associated with comprehension of complex social interaction and theory of mind. Our finding could be an example of diverging effects on social cognitive development between enthusiasm and assertiveness.

### Open practices statement

Data can be made available to researchers upon reasonable request. The study was not preregistered.

## Acknowledgements

This research was supported by grants from the Swedish Research Council (2014-01160, 2018-01322) and The Bank of Sweden Tercentenary Foundation (P20-0125).

## Notes

### Competing Interest Statement

The authors have declared no competing interest.

